# Impact of sequence variant detection and bacterial DNA extraction methods on the measurement of microbial community composition in human stool

**DOI:** 10.1101/212134

**Authors:** Riley Hughes, Zeynep Alkan, Nancy L. Keim, Mary E. Kable

**Author notes:** **Corresponding Author:** Mary E. Kable.

## Abstract

**Background:** The human gut microbiome has been widely studied in the context of human health and metabolism, however the question of how to analyze this community remains contentious. This study compares new and previously well established methods aimed at reducing bias in bioinformatics analysis (QIIME 1 and DADA2) and bacterial DNA extraction of human fecal samples in 16S rRNA marker gene surveys.

**Results:** Analysis of a mock DNA community using DADA2 identified more chimeras (QIIME 1: 0.70% of total reads vs DADA2: 1.96%), fewer sequence variants, (QIIME 1: 1297.4 + 98.88 vs. DADA2: 136.27 + 11.35, mean + SD) and correct taxa at a higher resolution of classification (i.e. genus-level) than open reference OTU picking in QIIME 1. Additionally, the extraction of whole cell mock community bacterial DNA using four commercially available kits resulted in varying DNA yield, quality and bacterial community composition. Of the four kits compared, ZymoBIOMICS DNA Miniprep Kit provided the greatest yield, with a slight enrichment of *Enterococcus.* However, QIAamp Fast DNA Stool Mini Kit resulted in the highest DNA quality. Mo Bio PowerFecal DNA Kit had the most dramatic effect on the mock community composition, resulting in an increased proportion of members of the family *Enterobacteriaceae* and genus *Eshcerichia* as well as members of genera *Lactobacillus* and *Pseudomonas.* The presence of a sterile fecal matrix had a slight, but inconsistent effect on the yield, quality and taxa identified after extraction with all four DNA extraction kits. Extraction of bacterial DNA from native stool samples revealed a distinct effect of the DNA stabilization reagent DNA/RNA Shield on community composition, causing an increase in the detected abundance of members of orders *Bifidobacteriales*, *Bacteroidales*, *Turicibacterales*, *Clostridiales* and *Enterobacteriales*.

**Conclusion:** These results confirm that the DADA2 algorithm is superior to sequence clustering by similarity to determine microbial community structure. Additionally, commercially available kits used for bacterial DNA extraction from fecal samples have some effect on the proportion of high abundance members detected in a microbial community, but it is less significant than the effect of using DNA stabilization reagent, DNA/RNA Shield.

## INTRODUCTION

Marker gene surveys utilizing PCR amplification of a short region of the bacterial 16S rRNA gene from bacterial DNA extracted from environmental samples are becoming increasingly affordable, leading to their ubiquitous implementation in nearly every aspect of biological sciences research [1-8]. However, this method can be heavily affected by technical bias, which is induced at each step in the experimental protocol required to generate marker gene data including; sample handling, bacterial DNA extraction, PCR amplification, sequencing and bioinformatics anlysis [9, 10]. PCR conditions and primer choice can impact biases during the amplification process, which has downstream effects on library preparation and formation of chimeric sequences [11, 12]. However, the two most important sources of technical bias, which can be relatively easily controlled, in marker gene surveys are DNA extraction and bioinformatics analysis [13].

Clustering into operational taxonomic units (OTUs) has been one of the primary bioinformatic methods used to group and identify bacterial taxa in samples in metagenomics and marker gene based sequencing analyses. This method utilizes percent sequence similarity to group sequences into operational taxonomic units (OTUs). The common similarity threshold used to define these OTUs is 97%, which is based on a study showing that most strains have 97% 16S rRNA sequence similarity [14]. From each OTU cluster, a single sequence is selected as the “representative sequence” and is classified based on a reference database. All sequences within the OTU cluster are then given the same taxonomic classification. OTU clustering offers a computational benefit, reducing millions of reads into only thousands of OTUs, allowing for rapid analysis of datasets [15].

However, the OTU clustering method has long been understood to have a number of drawbacks [15]. For instance, percent sequence similarity can overestimate the evolutionary similarity between sequences, leading to inappropriate clustering of sequences. Additionally, the standard 97% sequence similarity used to define species is an approximation and varies between taxa [16]. Higher rate of false positives (i.e. identification of taxa not present in the sample) as well as poor sequence and taxonomic resolution have also been cited as issues with OTU clustering [17, 18]. With the development of a number of new algorithms for sequence variant identification including Devisive Amplicon Denoising Algorithm (DADA2), unoise2, minimum entropy decomposition and Deblur [19-23], additional criticisms have surfaced regarding the OTU clustering method [24, 25] and the need to conduct and publish independent direct comparisons of methods has arisen.

Before sequences can even be analyzed and results affected by OTU clustering vs. sequence variant detection, DNA extraction methods can heavily influence the proportion of bacterial taxa detected in an environmental sample. Previous studies investigating the impact of various DNA extraction methods on 16S rRNA analyses of stool microbial communities each lack the combined use of a mock community in the relevant stool matrix background.

Additionally, the number of optimizable steps in DNA extraction protocols results in a near infinte number of possible ways to execute this type of experiment. Most notably, previous efforts to compare DNA extraction methods have indicated that the bead beating protocol tends to be the source of greatest variation between kits [26-28], yet few if any have held this variable constant during comparison. Finally, as technology evolves, new DNA extraction kits and bioinformatics methods are constantly being developed. Therefore, the need to compare and analyze new methods remains.

In this study we perform two important comparisons. First, we examine DADA2’s core denoising algorithm relative to the open reference OTU clustering method used in QIIME 1 to confirm which method results in a more accurate classification of the taxa present in a predefined mock community of bacteria. Second, we use a whole cell mock community in a sterile feces background to compare four relevant DNA extraction methods [10, 13, 29] with standardized speed and duration of bead beating.

## METHODS

### Preparation of stool samples

Whole stool samples were collected at home by three human subjects, placed in a cooler containing ice and brought to the Western Human Nutrition Research Center within 12 h of generation. Upon arrival at the facility, each sample was stored briefly at 4ºC until it could be homogenized in a stomacher for three minutes and flash frozen on dry ice. These samples were thawed, combined in equal amounts, mixed, and then divided into 2 pools. The first pool, which will be referred to as “native stool”, contained 1 g of stool from each subject, combined by homogenization in a stomacher twice for 5 min. A portion of this 3 g mixture was set aside in 100 mg aliquots for DNA extraction. The remaining 1 g of the mixture was combined with 9 mL of nucleotide stabilization reagent (DNA/RNA Shield, Zymo Research, Irvine, CA) by vortexing and incubated at room temperature overnight before 250 mg aliquots were weighed out for DNA extraction.

A “sterile” fecal sample was prepared as previously described [30] from the second pool, which contained 3 g of stool from each subject. Briefly, the 9 g mixture was stirred together with 90 mL of boiling 30% hydrogen peroxide (H_2_O_2_) for 15 min. The boiled stool mixture was then passed over a 0.22 µm vacuum filter (Sarstedt, Nümbrecht, Germany) to collect particulate matter. Fecal particulate matter retained on the filter was then washed with sterile phosphate buffered saline (DPBS, pH=7.0-7.3, ThermoFisher, Waltham, MA) in 100 mL batches until H_2_O_2_ was no longer detected in the filtrate using detection strips (MQuant Peroxide Test, MilliporeSigma, St. Louis, MO). This required 1 L of PBS. A total of 3.1 g of dry particulate matter was collected from the filter surface and suspended in 4.5 mL sterile PBS to create a sterile fecal matix. To create a mock stool sample with a known bacterial community, a 1.5g aliquot of this sterile feces was homogenized in a stomacher for 2 min together with 1.125 mL of commercially available whole cell mock community (ZymoBIOMICS Microbial Community Standard, Zymo Research, Irvine, CA, lot number ZRC183430) and this mixture was set aside in 173 mg aliquots for DNA extraction (75 µL of mock community per 100 mg of stool). The microbial strains included in the standard along with their theoretical relative abundances are listed in Table S1. The remainder of the sterile fecal matrix was portioned into 100 mg aliquots as blank controls for DNA extraction.

### Experimental design and bacterial DNA extraction

A total of six sample types were prepared for DNA extraction; (1) kit blank with no sample, (2) 75 µL mock community alone, (3) 100 mg sterile feces alone, (4) sterile feces with mock community added totaling 173 mg as described above, (5) 100 mg native stool and (6) 25 mg native stool suspended in nucleotide stabilization reagent (DNA/RNA Shield, Zymo Research, Irvine, CA) totaling 250 mg. Twelve aliquots of each sample type (three per kit) were homogenized 5 times in bead tubes from three of the DNA extraction kits or bead tubes prepared separately (described below) at 6.5 m/s for 1 min using a homogenizer (FastPrep-24 Classic Instrument, MP Biomedicals). Samples were rested on ice for three minutes between each shaking interval. Bacterial DNA was then extracted using (1) QIAamp Fast DNA Stool Mini Kit, (2) MO BIO PowerFecal DNA Kit, (3) ZR Fecal DNA Kit and (4) ZymoBIOMICS DNA Miniprep Kit. For the QIAamp Fast DNA Stool Mini Kit, which contains no bead tubes, sterile 2 mL screw cap tubes containing 300 mg of 0.1 mm dimeter zirconia/silica beads (BioSpec Products, Bartlesville, OK) were prepared separately and sterilized by autoclaving. After homogenization by bead beating, the manufacturer’s protocol was followed for each kit with the following exceptions:

*All kits* - Wash and elution buffers were incubated on the column for 10 min prior to centrifugation. Before the addition of elution buffer, columns were centrifuged for three minutes with caps open in order to completely remove wash buffers.

*QIAamp Fast DNA Stool Mini Kit –* The protocol for “Isolation of DNA from Stool for Pathogen Detection” in the QIAamp Fast DNA Stool Mini Handbook (03/2014) was used with few modifications. Briefly, after bead-beating stool samples in 1 mL InhibitEx buffer, the samples were heated at 95ºC for 5 minutes. Centrifugation to remove particulate matter was performed for 3 min on the whole sample and for an additional 3 min on the resulting supernatant. A larger portion than recommended, 400 µL, of the clarified sample was transferred to a new tube containing 30 µL proteinase K. Additional lysis was performed as described in the manufacterer’s protocol. However, only 200 µL of lysate was added to the QIAamp spin column. DNA was eluted with 30 µL buffer ATE.

*MO BIO (now QIAamp) PowerFecal DNA Kit –*DNA was eluted with 50 µL buffer C6.

*ZR Fecal DNA Kit (now Quick-DNA Fecal/Soil Microbe Miniprep Kit) –* DNA was eluted in 50 µL DNAse free water.

*ZymoBIOMICS DNA Miniprep Kit –* DNA was eluted in 50 µL DNAse free water.

### Amplification and sequencing of 16S rRNA

The 16S rRNA V4 region was amplified as previously described [31] using primers F515 and R806 [32]. A unique eight nucleotide Hamming code sequence was included on the 5’ end of F515 [33, 34] for amplification of each sample. Each 50 µL reaction mixture was composed of 20 ng of template DNA, 1.5 U Ex Taq DNA polymerase (TaKaRa, Otsu, Japan), 100 nM of forward primer, 100 nM reverse primer, 500 nM magnesium chloride, 200 nM dNTPs and 1X Ex Taq buffer. Amplification was performed in triplicate for each sample with one cycle at 94°C for 3 min followed by 25 cycles of 94°C for 45 s, 50°C for 60 s, and 72°C for 60 s. A final extension step was performed at 72°C for 10 min. Equal volumes of each PCR reaction (40 µL) were pooled and gel purified with the Wizard SV Gel and PCR cleanup system (Promega, Madison, WI). Ligation of NEXTflex adapters (Bioo Scientific, Austin, TX) and 300-bp paired end sequencing on an Illumina MiSeq instrument with MiSeq Reagent Kit v3 (Illumina) was performed at the University of California, Davis (http://dnatech.genomecenter.ucdavis.edu/).

In order to eliminate the bias introduced by PCR amplification and sequencing from our downstream analyses, a commercially available mock microbial community DNA standard (ZymoBIOMICS™ Microbial Community DNA Standard, lot number ZRC187324), a sample which we will refer to as Mock DNA was amplified and sequenced in the same manner as all other experimental samples. The DNA standard is a mixture of genomic DNA extracted and quantified from pure cultures of eight bacterial and two fungal strains with the same theoretical composition as the whole cell mock community described above. Metagenomic sequencing, was performed by Zymo Research as part of their product quality assesment to determine the percent relative abundance of the microbial strains in both the DNA and whole cell standards. Their reported results are listed in Table S1.

### 16S rRNA gene sequence analysis

A summary of the methods used for analysis is described in Table 1. FASTQ files were analyzed using QIIME version 1.9.1 [35], which will hereafter be referred to as QIIME 1, or DADA2 version 1.4.0 [20]. R version 3.4.0 was used for all analyses. For the QIIME 1 analysis, referred to throughout this manuscript, barcodes were extracted and the split_libraries_fastq.py script was used for demultiplexing and quality filtering. Demultiplexing was performed only with barcodes containing no sequencing errors, and quality filtering was performed at a Phred quality threshold of 29. Chimeric sequences were identified with identify_chimeric_seqs.py using usearch [36] and removed. The remaining DNA sequences were grouped into OTUs with 97% matched sequence identity by the use of pick_open_reference_otus.py. The default for open reference OTU picking in QIIME is to use the first read as the representative sequence to form the OTU clusters. In order to more closely imitate the DADA2 pipeline, this default behavior was changed to use the most abundant sequence by passing a parameter file using the function pick_rep_set.py (method most_abundant). Otherwise default parameters were used. Greengenes 13.8 was used as the reference database [37] for chimera checking, OTU picking, and taxonomy assignment.

**Table 1.** Summary of bioinformatic methods

| Step | DADA2 1.4.0 | QIIME 1.9.1 |
| --- | --- | --- |
| Input | demultiplexed fastq files + mapping file | fastq files + mapping file |
| Pre-Processing | - Filter and trim (truncLen=190, otherwise standard parameters)<br>- Dereplication | - Extract barcodes and remove primers<br>- Split libraries (demultiplex and quality filter – Q=29, otherwise default parameters) |
| Pick OTUs/variants | Sequence-variant inference (Sample inference/Denoising) | Open reference OTU picking (usearch, pick_rep_set method most_abundant, otherwise default parameters) |
| Remove chimeras | Remove bimeras | Remove chimeras* (usearch) |
| Assign taxonomy | Greengenes v13_8_99 | Greengenes v13_8_99 |

\*Chimera/bimera removal comes before OTU picking in QIIME 1 but after Sample Inference in DADA2

DADA2’s denoising algorithm is based on pairwise comparison of sequences and uses quality scores of the reads as well as the probability of various copy errors (transition probabilities) that could be introduced during replication and sequencing. See Callahan et al. [20] for full documentation of the core DADA2 algorithm. Methods used for DADA2 analyisis were adapted from the DADA2 Pipline Tutorial (1.4) and DADA2 Frequently Asked Questions, which are both currently available in the DADA2 GitHub documentation. Brielfly, prior to analyses in DADA2, samples were demultiplexed using the QIIME 1.9.1 split_libraries_fastq.py script with the following modifications from default parameters: -r (max bad run length) 999, -n (max length of sequence) 999, -q (Phred quality threshold) 0, -p (min number of high quality bases as fraction of read length) 0.0001 and --store_demultiplexed_fastq. This removed the majority of quality filtering that is typically implemented by the QIIME 1 pipeline using this script and created individual fastq files for each sample. The demultiplexed files were used as the input for DADA2.

Quality profiles of the reads were analyzed using the DADA2 function, plotQualityProfile, to determine positions at which read quality greatly diminished. Reads were then filtered and trimmed at the identified positions (truncLen=190) using the filterAndTrim function with standard parameters (maxN=0, truncQ=2,and maxEE=2). Dereplication was then used to identify all unique sequences present in the data set and determine the abundance of each sequence. DADA2 also retains a summary of the quality information associated with each unique sequence, using this to inform the error model of the subsequent denoising step, increasing its accuracy [20]. DADA2’s error model automatically filters out singletons, removing them before the subsequent sample inference step. Quality of the error estimation was then visualized using the plotErrors function to ensure good fit. Sample inference was performed using the inferred error model and chimeric sequences were removed using the removeBimeraDenovo function. It is relevant to note that DADA2 implements bimera removal after sample inference has been performed, whereas removal of chimeras in QIIME 1 occurs before the OTU picking step. The Greengenes 13.8 database was used to assign taxonomy using the assignTaxonomy function.

### Statistical Analysis

OTU or sequence variant counts and rarefaction curves were determined on sequence count files (referred to as sequence table and OTU table in DADA2 and QIIME 1 respectively) generated by each analysis pipeline. These were determined using a count of the number of rows in each output file that contained non-zero values, referred to as non-zero OTU/SV counts, for each sample.

Analysis of the relative proportion of each bacterial taxa was made after the data were rarefied at a sequencing depth of 50,000 sequences per sample for both QIIME 1 and DADA2. The rarefied sequence variant counts were summed by taxonomic identification and differential abundances between experimental groups were determined using LefSe [38]. This method involves the Kruskal-Wallis (KW) sum-rank test between classes of data followed by (unpaired) Wilcoxon rank-sum test to conduct pairwise tests among subclasses. LDA is then used to estimate the effect size for each of the identified taxa. We used LEfSe (Galaxy Version 1.0) with default paramters (α KW = 0.05; α Wilcoxon = 0.05; LDA score threshold = 2.0) as well as using the all-against-all strategy for multi-class analysis. All other comparisons were made using either Welch’s t-test or Kruskal-Wallis (KW).

## RESULTS

### The DADA2 denoising algorithm improves accuracy of bacterial community measurement

QIIME 1 and DADA2 were compared using 18,651,434 sequences generated by Illumina MiSeq sequencing of 6 individual PCR amplifications of a microbial community DNA standard (Mock DNA). After demultiplexing and quality filtering using QIIME 1, 790,502 total sequences remained. Of these, 5,532 chimeras were identified using usearch, accounting for only 0.70% of total sequences. On the other hand, the trimming, denoising and dereplication steps of DADA2 resulted in 368 sequences (or inferred variants), which could be considered more equivalent to a representative set of sequences picked by open reference OTU picking. Out of these sequences, 160 bimeras were identified, representing 43.48% of inferred variants, but only 1.96% of total reads after dereplication, and filtering (1,354,268 reads), which is still nearly double the percentage detected using usearch in QIIME 1.

QIIME 1 identified a much larger number of OTUs/SVs than DADA2 in Mock DNA (QIIME 1: 1145.5 + 68.73 vs. DADA2: 123.5 + 8.12, mean + SD) (**Figure 1A**). However, DADA2 still greatly overestimated the number of non-zero variants relative to the expected number of bacterial species present in the Mock DNA samples. Low abundance sequences identified by DADA2 were investigated further. The Hamming distance of low abundant sequences relative to more abundant sequence-variants they were split from fell in a range from 1 to 80, and quality scores at nucleotide positions used to determine a particular low abundance sequence was unique relative to the more abundant sequence it was split away from were above 29. However, when BLAST was used to compare these low abundance sequences to those available in the National Center for Biotechnology Information (NCBI) nucleotide database, 87% of uniqe sequences in the Mock DNA samples were exact matches (100% query cover, 100% identity) to bacterial taxa that tend to be abundant in human stool samples, such as genera *Bifidobacterium, Turicibacter, and Blautia*.

**Figure 1.**
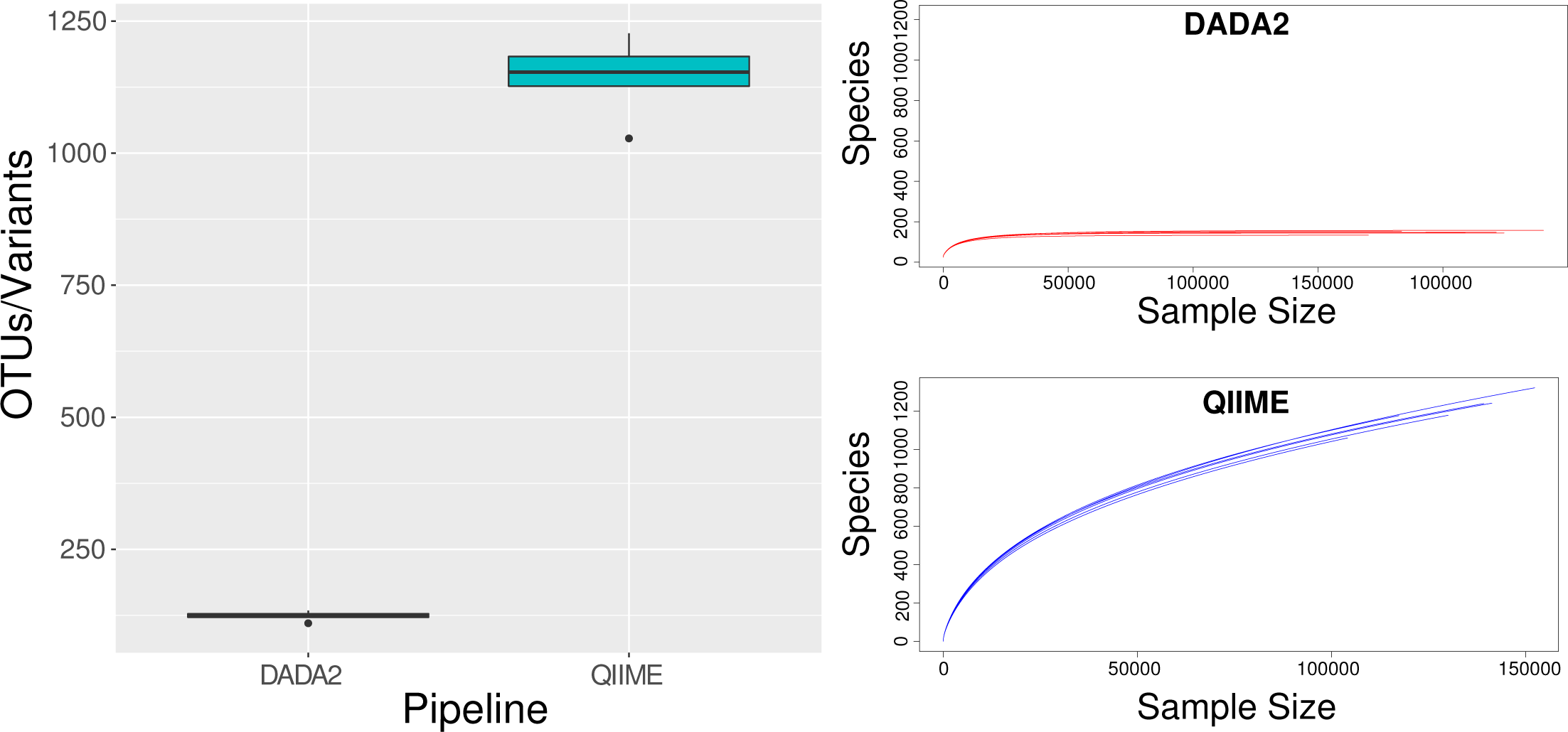
Variant counts resulting from QIIME 1 and DADA2 analyses. A) Boxplot of comparison between DADA2 and QIIME 1 OTU/sequence variant counts for Mock DNA. B) Rarefaction curves showing differences in taxonomic discovery rate between DADA2 and QIIME 1. Six individually amplified and sequenced Mock DNA samples were analyzed with each pipeline.

Rarefaction curves representing the discovery rate of unique sequences, potentially attributed to new taxonomic units, as a function of sequencing effort (i.e. number of sequences) [39], reflected the differences in non-zero OTU/SV counts between QIIME 1 and DADA2 (**Figure 1B and C**). As sequencing effort increases, QIIME 1 open reference OTU clustering results in the detection of continually increasing numbers of unique sequences in Mock DNA samples. However, the number of unique sequences detected by DADA2 does not increase with sequencing effort in the same way as for QIIME 1, instead the number of unique sequences detected levels out at approximately 50,000 sequences per sample.

While QIIME 1 identified many more OTUs/SVs than DADA2, rarefaction at 50,000 sequences per sample followed by removal of low abundance taxa (<1%) into a category termed “Other”, showed a similar taxonomic profile of the Mock DNA samples detected by both QIIME 1 and DADA2 (**Figure 2**). However, DADA2 identified correct taxa at a higher resolution of classification (i.e. genus-level) with less redundancy (i.e. identification of the same taxa at different levels of taxonomic classification, such as f_*Bacillaceae* and g_*Bacillus*) than QIIME 1. More specifically, eight taxa were present at greater than 1% relative abundance as detected by DADA2. Seven out of these eight were correctly identified to the genus level. The last variant was correctly identified at the family level (e.g. f_*Enterobacteriaceae* includes *Salmonella enterica*). QIIME 1 identified nine taxa present at greater than 1% relative abundance. Out of these nine, four were redundant at different levels of phylogenetic resolution. These included f__*Bacillaceae* and g__*Bacillus* as well as f__*Pseudomonadaceae* and g__*Pseudomonas*. All nine taxa identified were present in the Mock DNA community (no spurious identification), but two taxa remained classified only to the family level (f__*Enterobacteriaceae* and f__*Listeriaceae*) (**Figure 2**). LefSe analysis showed significant differences in the majority of taxa identified excluding only g__*Enterococcus* and g__*Staphylococcus*. Because of this increased accuracy in taxonomic identification, the remainder of comparisons examining DNA extraction kits were analyzed using DADA2.

**Figure 2.**
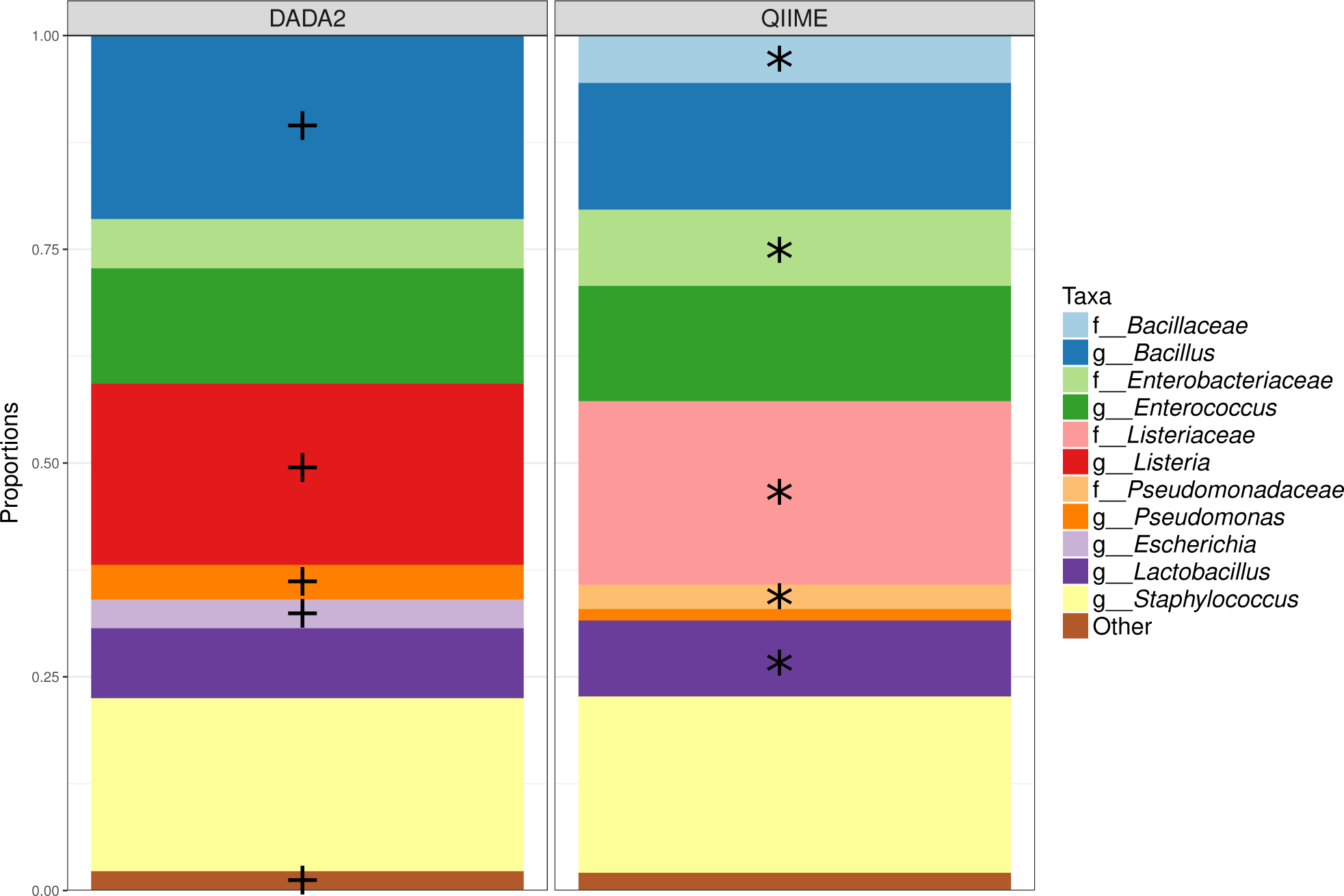
Relative taxonomic abundance of Mock Community DNA samples analyzed by DADA2 and QIIME 1. OTU and sequence variant counts were rarefied at 50,000 sequences per sample for both groups. All taxa present at <1% abundance were grouped into the “Other” category. Each bar represents six PCR amplifications of Mock DNA. +Signficantly enriched in DADA2 analyzed samples. *Significantly enriched in QIIME 1 analyzed samples.

### DNA yield and quality vary among extraction kits

The efficiency of four commercial DNA extraction kits was assessed using commercially available whole cell mock community (Mock Community) and the whole cell mock community spiked into sterilized fecal matrix (Mock Community in Sterilized Feces). There was a significant difference among the kits in DNA yield (KW Mock Community *P* = 0.02488, Mock Community in Sterilized Feces *P* = 0.01556) and quality (KW Mock Community p = 0.03781,

Mock Community in Sterilized Feces *P* = 0.04358) from both sample types. ZR Fecal and ZymoBIOMICS delivered the highest quantity of DNA for both the whole cell Mock Community alone (ZR Fecal average = 59.3 ng/uL, ZymoBIOMICS average = 58.8 ng/uL) and Mock Community in Sterile Feces (ZR Fecal average = 39.9ng/uL, ZymoBIOMICS average = 31.8 ng/uL). However, QIAamp delivered the highest quality DNA from both sample types (A260/A280 = 2.5 and 1.86 respectively) (**Figure. 3A and B**). The DNA yield and quality were also affected by the presence of the sterile feces matrix. Both decreased in the presence of the matrix for each kit, except for QIAamp. However, the difference in yield was only significant in the ZR Fecal (Welch’s t-test *P* = 0.02699) and ZymoBIOMICS (*P* = 0.008911) kits and the difference in quality was only significant in the ZR Fecal kit (*P* = 0.03097).

**Figure 3.**
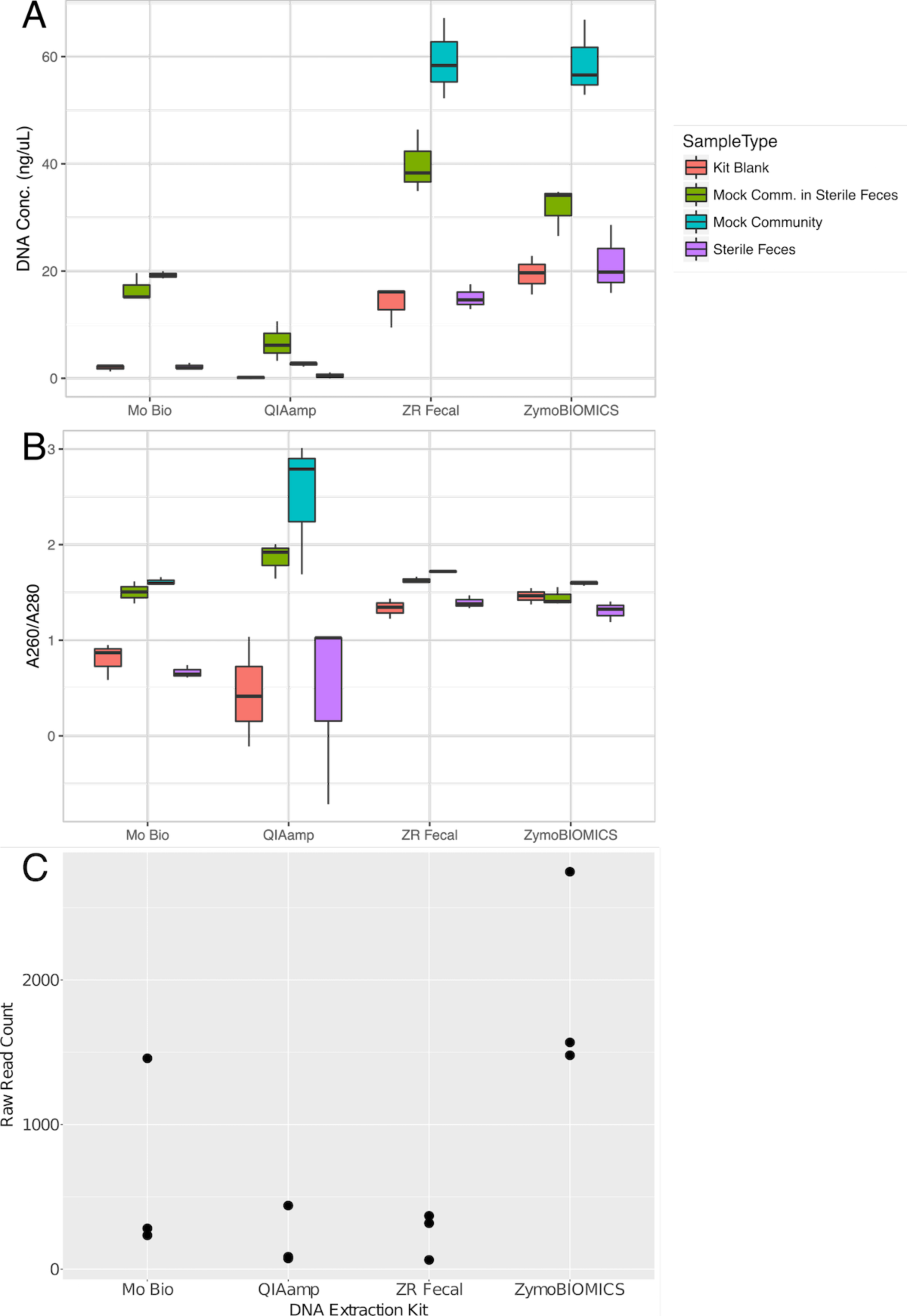
Yield and quality of mock community DNA extracted by four commercial kits. Boxplots showing A) DNA yield (ng/uL), B) DNA quality (A260/A280) and C) raw sequence counts obtained from whole cell mock community (Mock Community) or whole cell mock community spiked into a sterile feces matrix (Mock Comm in Sterile Feces), sterile feces alone (Sterile Feces) or no sample (Kit Blank) using each of four commercial DNA extraction kits (MoBio, Qiagen, ZRFecal and ZymoBIOMICS). Three of each sample type are represented.

The yield obtained from blank samples followed the same trend as the yield obtained from mock community samples. It was significantly higher for ZR Fecal and ZymoBIOMICS than it was for the other two protocols, reaching levels greater than 10 ng/uL for each of the two kits. However, the number of bacterial sequences detected after PCR and sequencing in the blanks were not significantly different among kits (**Figure 3C**, KW Blank p-value = 0.09234).

### Measurement of bacterial community composition is affected by DNA extraction protoco

In addition to DNA yield and quality, the proportion of bacterial taxa measured after extraction with each kit was determined. The relative proportions of taxa expected to most closely represent reality were determined using the Mock DNA standard described above.

Weighted UniFrac distances between extracted samples and the Mock DNA samples were visualized by principal componants analysis (**Figure 4A**) and summarized in boxplots (**Figure 4B**). Samples extracted with the Mo Bio kit had the greatest combined distance from Mock DNA (mean=0.0429, median=0.0409) compared to the other kits (Mo Bio mean = 0.0107, median = 0.0121; ZymoBIOMICS mean = 0.0039, median = 0.0034; ZR Fecal mean = 0.0097, median = 0.0078). Distances were significantly affected by the presence of a sterile fecal matrix in all kits examined (Mo Bio *P* =1.36e-05; QIAamp *P* < 2.2e-16; ZymoBIOMICS *P* = 1.887e-6; ZR Fecal *P* = 0.0131). In the case of the Mo Bio and ZR Fecal kits, the presence of a stool matrix decreased the distance from Mock DNA (Mo Bio mean with matrix = 0.0378, mean without matrix = 0.0479; ZR Fecal mean with matrix = 0.0078, mean without matrix = 0.0117).

**Figure 4.**
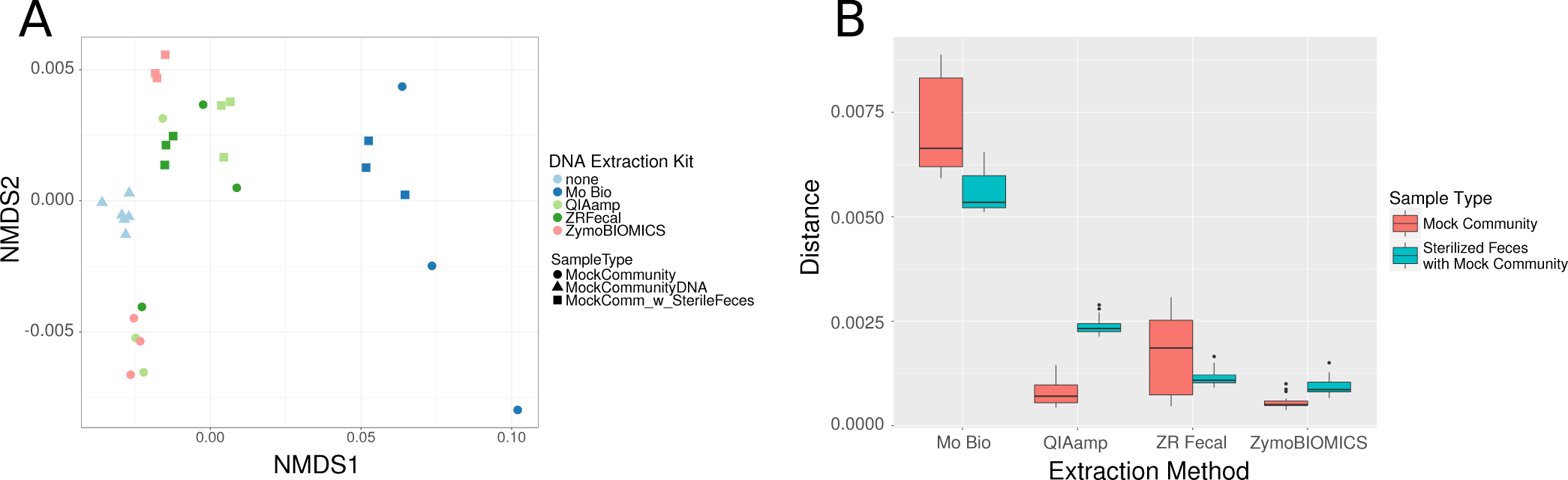
Weighted UniFrac of distance between extracted mock community samples and Mock DNA. A) Principal coordinate analysis of weighted UniFrac distances among Mock Community or Mock Community in Sterile Feces and Mock DNA (control) samples. B) Boxplots summarizing the weighted UniFrac distance between Mock DNA and each extracted sample type grouped by extraction kit (Mo Bio, QIAamp, ZR Fecal and ZymoBIOMICS).

However, the opposite phenomenon occurred for the Qiagen kit protocol (mean with matrix = 0.0162, mean without matrix = 0.0053) and the ZymoBIOMICS kit (mean with matrix = 0.0064, mean without matrix = 0.0039).

LEfSe analysis identified the greatest number of significantly different taxa in Mock community samples extracted with the Mo Bio kit. This included an increased proportion of members of the family *Enterobacteriaceae* and genus *Eshcerichia* as well as members of genera *Lactobacillus* and *Pseudomonas* (Figure. 5). Mo Bio also enriched the “Other” category, indicating enrichment in several other low abundance taxa. Relative to the Mock DNA, decreased abundance of members of the phylum *Firmicutes*, including order *Bacillales* and class *Bacilli* and genus *Listeria,* though not genus *Bacillus* were detected in all extracted samples.

**Figure 5.**
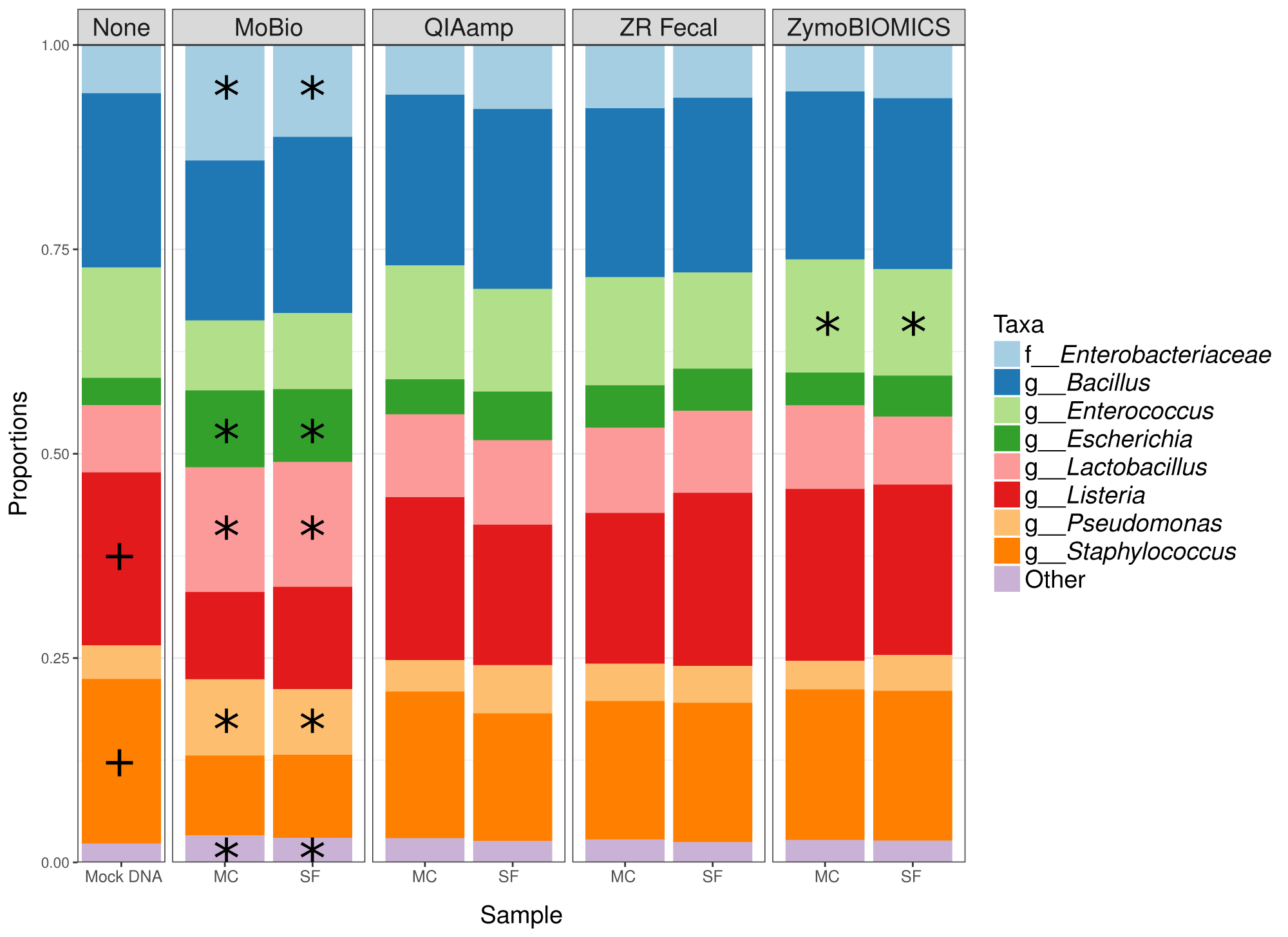
Relative taxonomic abundance of mock community taxa after extraction by four commercial kits. The proportions of taxa present in the Mock DNA sample are shown for comparison (None). Each bar represents a summary of technical replicates (six Mock DNA samples and three of each of the other sample types). MC is used to designate whole cell mock community only and SF is used to designate mock community spiked into sterile feces matrix. +Signficantly enriched in Mock DNA samples. *Significantly enriched in extracted samples.

Members of the gram positive genus *Staphylococcus* were also proportionally decreased in the extracted samples relative to Mock DNA. Mock community samples extracted by ZymoBIOMICS showed significant enrichment of genus *Enterococcus*.

### Use of nucleotide stabilization reagent significantly affects measurement of microbial community composition

After assessing the performance of different pipelines and extraction kits on the mock community, we looked to confirm the relative efficiency of each kit and further investigate the effect of the nucleotide stabilization reagent, DNA/RNA Shield, using a representative pool of natural or native stool samples (Native Stool and Native Stool with DNA/RNA Shield). DNA yield was significantly different among kits for extraction from pooled native stool samples similar to observations for the mock community samples above (KW p-value = 0.0329 native stool; 0.01556 native stool in shield). Additionally, the presence of stabilization reagent affected the amount of DNA recovered by each kit. For both kits from Zymo Research (ZR Fecal and ZymoBIOMICS), the amount of DNA recovered per gram of stool was significantly increased (p-value = 0.0002916 and 0.01315) in the presence of stabilization reagent (**Figure 6A**). This was not true for the other two protocols which showed a decrease. Although, the decrease was only significant for the QIAamp kit (p-value = 0.003795). The quality of DNA recovered was also significantly different among kits for extraction of the Native Stool and Native Stool with DNA/RNA Shield (KW p-value = 0.02871; 0.01879), with QIAamp again providing the highest quality DNA (**Figure 6B**). However, the quality of DNA was only significantly affected by the presence of DNA/RNA Shield during extraction with the Mo Bio PowerFecal Kit (p-value = 6.435e-05).

**Figure 6.**
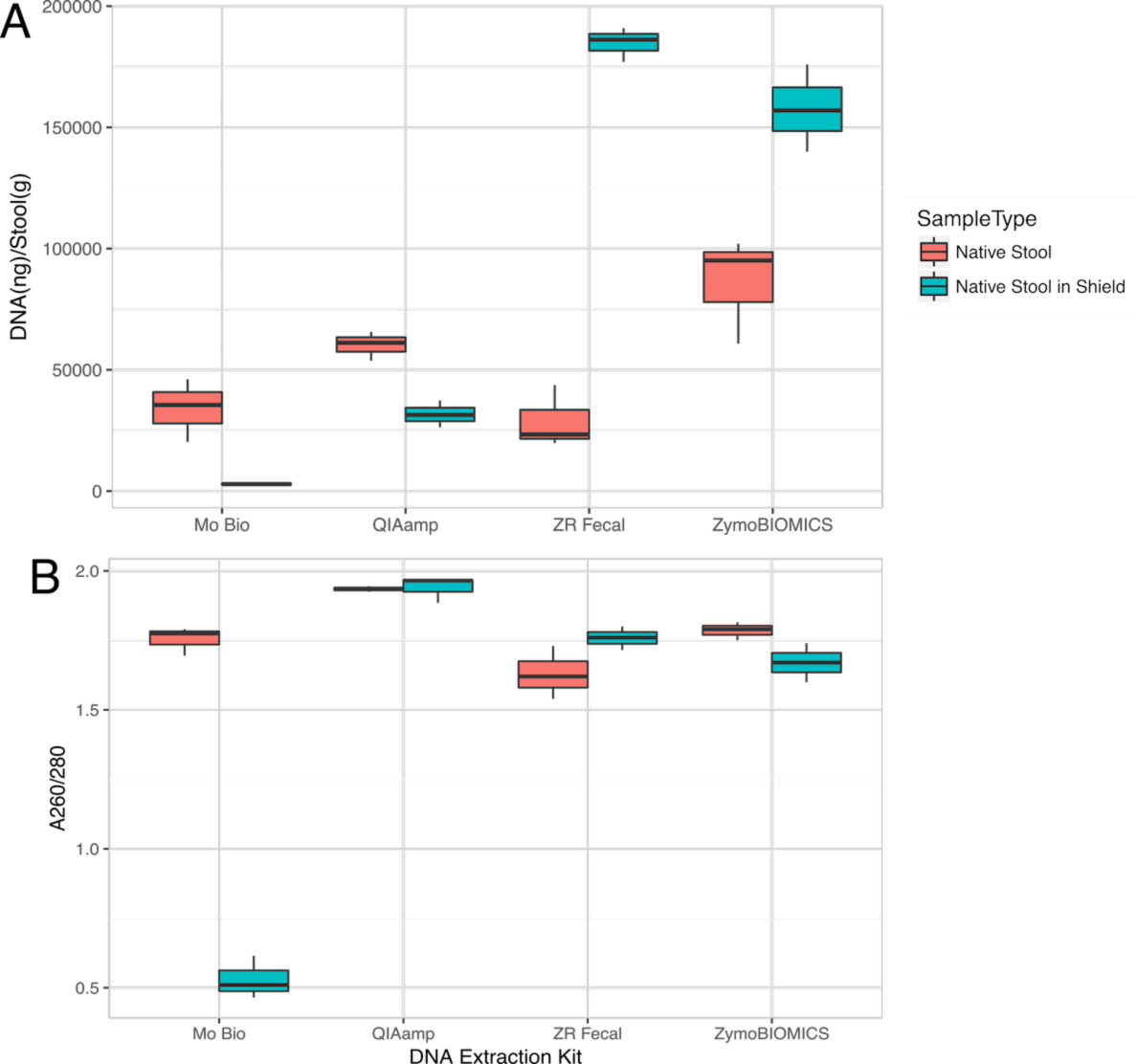
Yield and quality of pooled native stool bacterial community DNA extracted by four commercial kits. Boxplots showing A) DNA yield in ng/g of stool and B) quality (A260/A280) obtained from pooled (three stool samples) native stool community (Native Stool) and pooled native stool community suspended in nucleotide stabilization reagent (Native Stool in DNA Shield) using each of four commercial DNA extraction kits (Mo Bio, QIAamp, ZR Fecal and ZymoBIOMICS). Three of each sample type are represented.

Principal coordinate analysis of weighted UniFrac distances showed that samples clustered by stabilization reagent first and by DNA extraction method second (**Figure 7A and B**). The impact of stabilization reagent on community composition was again greatest for the Mo Bio kit (**Figure 7C**). However, analysis of the relative abundance of bacterial taxa present after extraction with each kit showed significant differences in relative proportion of taxa enriched between samples with and without DNA/RNA Shield across all extraction kits (**Figure. 8**) Order *Clostridiales* including family *Ruminococcaceae* and genera *Clostridium, Oscillospira, Ruminococcus,* and *SMB53* as well as order *Bifidobacteriales* including genus *Bifidobacterium* were significantly increased in native stool with DNA/RNA Shield. Order *Bacteroidales* including families *Rikenellaceae* and *Porphyromonadaceae* and genera *Bacteroides* and *Parabacteroides*; order *Turicibacterales* including genus *Turicibacter*; and order *Enterobacteriales* including genus *Escherichia* were also enriched in samples with DNA/RNA Shield. While not significant at the order level, other members of *Firmicutes* and *Actinobacteria*, including genera *Dorea*, *Faecalibacterium*, *Eggerthella*, *Roseburia*, *Collinsella*, *Coprococcus*, and *Blautia* were decreased in the presence of stabilization reagent.

**Figure 7.**
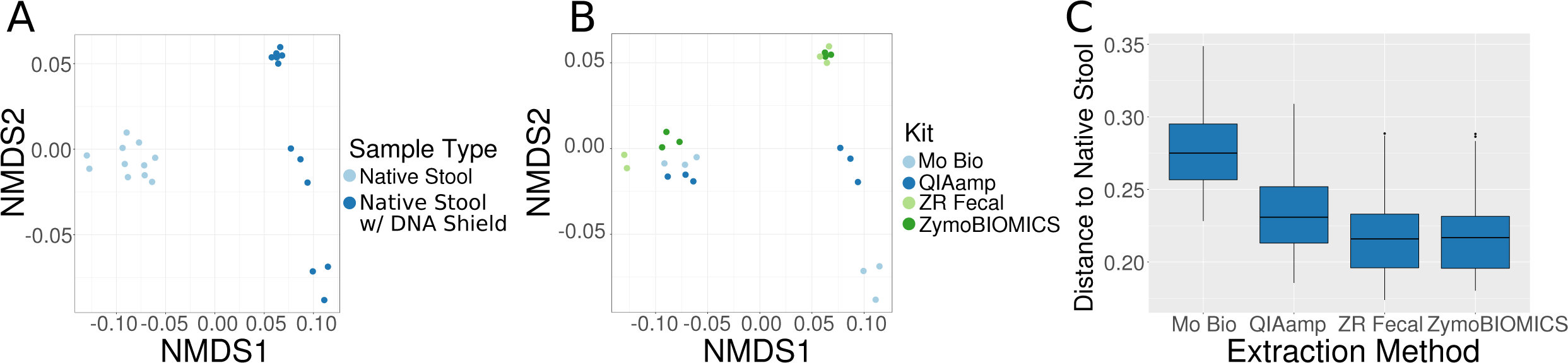
Relative composition of microbial communities in pooled native stool samples with and without stabilization reagent. Weighted UniFrac of distance between native stool samples colored by A) Sample Type with (dark blue) or without (light blue) DNA/RNA Shield B) Extraction Kit and C) Boxplot of the weighted UniFrac distance between Native Stool w/ DNA Shield samples and Native Stool w/o DNA Shield samples, separated by DNA Extraction Kit.

**Figure 8.**
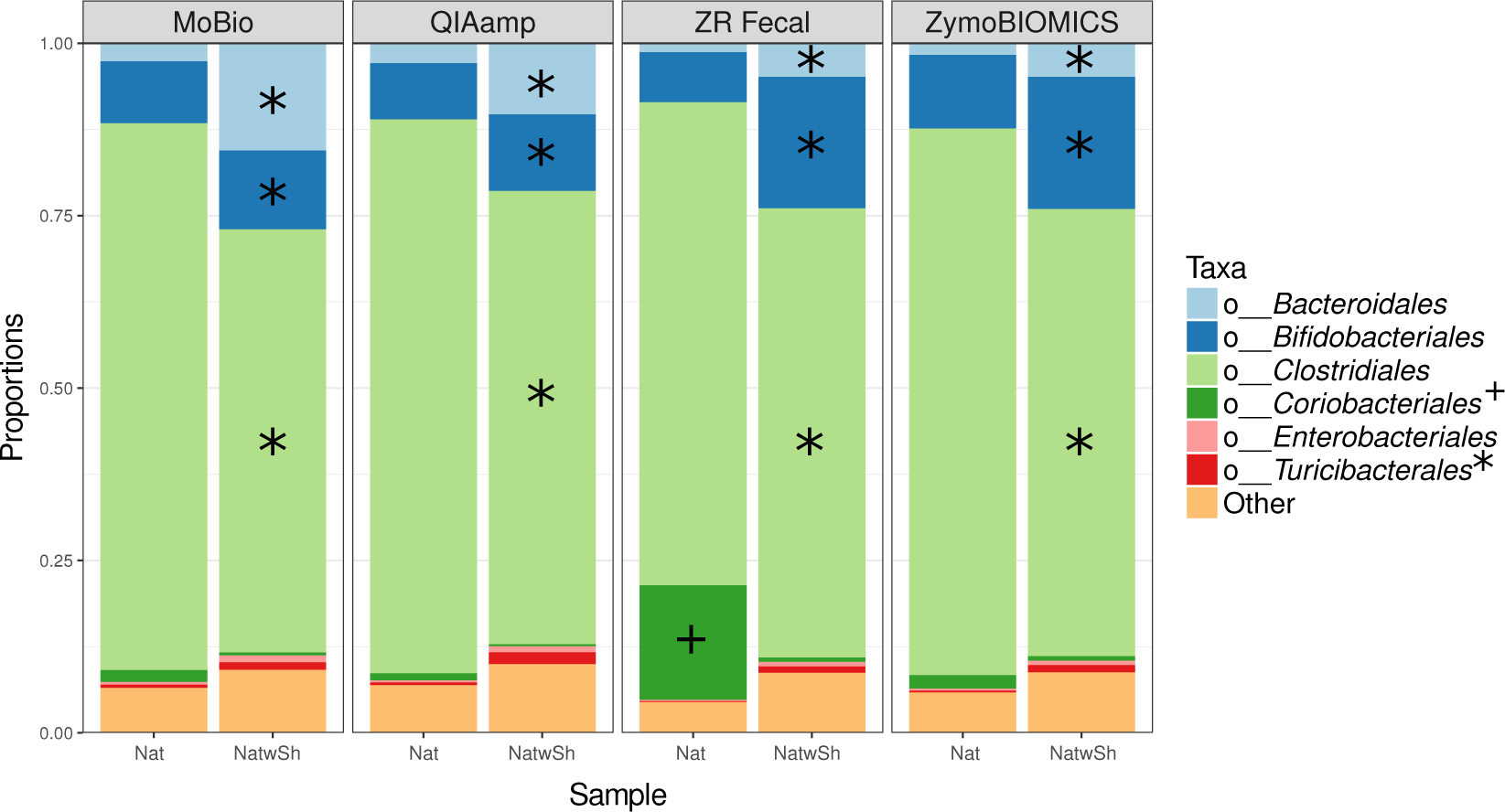
Relative abundance of taxa in pooled native stool. Relative proportions of taxa are shown at the order level for pooled native stool samples with (NatwSh) or without (Nat) pre-incubation in nucleotide stabilization reagent for each extraction kit. *Significantly enriched in samples without DNA shield. +Significantly enriched in samples with DNA shield.

## DISCUSSION

The determination of microbial community structure composition in environmental samples can be heavily affected by technical bias. As new methods are developed to deal with errors induced by DNA extraction, sequencing and other analysis methods, it remains necessary to empirically compare and validate each method using microbial standards. Here we have shown that DADA2 provides a more accurate assessment of the microbial community both in terms of the number of sequence-variants detected as well as the identity and phylogenetic resolution of taxa present. Additionally, if bead beating speed and duration are held constant, the commercially available kit used for bacterial DNA extraction from fecal samples has minimal effects on the proportion of high abundance members detected in a microbial community, except in the case of chemical incompatibility, which may be present between the Mo Bio kit and the DNA stabilization reagent, DNA/RNA Sheild.

The reduced number of unique sequences, identifiable to a higher taxonomic resolution detected using DADA2 relative to the QIIME 1 OTU clustering method was likely due to the method of error detection employed by DADA2, which statistically determines the most likely sequencing errors in a particular data set and then adjusts for them rather than rounding out by an allowable percent error (typically 97%). However, a number of low abundant taxa were also identified using DADA2 that were not present in the reference sequences for the mock community used for analysis. It should be noted that these taxa were detected without a stringint quality filter settting applied to the filterAndTrim function in DADA2. Therefore, it is possible that their number could be reduced further with a more stringint quality filter setting.

Optimization parameters aside, many of these taxa were abundant in DNA that was extracted from native stool samples at the same time as the mock community samples in this study. This indicates that some contamination of the Mock DNA sample occurred leading to a slightly greater number of detected sequence variants than we expected. However, because the same samples were analyzed using both pipelines, our conculsions regarding the improved accuracy of DADA2 remain valid.

Subsequent to the selection of a bioinformatics pipeline for our analyses, we found that DNA yield and quality varied among mock community samples and blanks from four commercially available DNA extraction kits. Given that the number of bacterial sequences detected in Zymo Research blanks were not significantly higher than in the other kits, it is unclear why the DNA yield was high in blank samples extracted using these two kits. We suppose that either, the chemistry involved in the Zymo Research kits results in absorbance at A260, or that there is viral or fungal DNA contaminant in the kit, which was undetected by our PCR protocol.

Within all four, both yield and quality were slightly impacted by the presence of sterile fecal matrix. The trend for a reduction in yield in the presence of matrix in three out of the four kits suggests that, as expected, the presence of physical impediments to bead-beating that tend to be present in the stool matrix, primarily undigested food particles, likely inhibit the effectiveness of the beads in disrupting bacterial cells [41, 42]. One exception was the QIAamp kit. However, the composition of the microbial community in the context of sterile fecal matrix was more dissimilar than mock community alone from the proportions predicted by our control. This did not result in a significantly detectable change in the relative proportions of abundant taxa, but insignificant increases in gram negative organisms and decreases in proportions of some gram positive organisms were observed. This would be expected if decreased efficiency of bacterial cell wall disruption by bead-beating occurred in the presence of the sterile fecal matrix. The Mo Bio kit, on the other hand, displayed decreased yield in the presence of sterile fecal matrix, but the microbial community composition tended to become more similar to the control than mock community along. The garnet beads included in the Mo Bio kit were pulverized at the speed and duration of shaking used in our protocol (see materials and methods). It is therefore possible that the very small broken particles of these beads disrupted bacterial cells so effectively that exposed DNA was also pulverized and the presence of a fecal matrix helped prevent some of this disruption.

Given that speed and duration of bead beating were held constant, the trends described for yield and quality across the four extraction kits in the presence of sterile fecal matrix suggest that the size, shape and composition of beads play role in the ability to sufficiently disrupt the stool matrix and facilitate the detection of “realistic” proportions of bacterial taxa. A second explanation for varying results across the kits, predominated by a slight decrease in nucleotide quality, in the presence of the sterile fecal matrix might be PCR inhibitors, such as carbohydrates, coming from the stool matrix, which are eliminated to differing degrees of completeness by each kit and could also be affected by use of stabilization or preservation reagent [43].

All four DNA extraction protocols showed a decreased relative abundance of the phylum *Firmicutes* and genus *Staphylococcus* extracted from whole cell mock community relative to the Mock DNA control sample. This may indicate that even the robust bead beating protocol used in this study (see Methods) was not sufficient to fully lyse all gram positive organisms contained in the stool samples. However, as shown in Table S1, the relative abundances of taxa in the whole cell mock community, as estimated by Zymo Research using metagenomics sequencing, differed from that in the Mock DNA. It is possible that the difference, reported by Zymo, was caused by biases introduced by the DNA extraction kit that they used to determine the abundance of their own community. Therefore, we are unable to use the difference between the “measured” values as the expected difference between the Mock DNA and mock microbial community samples as this would simply be a comparison of their extraction and sequencing methods and our own.

Given that both sample types were prepared with the same theoretical proportions, our analysis presumes that the Mock DNA is a close representation of the proportions in the whole cell mock community. Under this assumption, the ZymoBIOMICS DNA Miniprep Kit was determined to provide the closest representation of the “true” microbial community in a stool sample. On the other hand, the Mo Bio kit had the most distinct deviation from the expected microbial community composition. In the mock microbial community, characterized by significant increases in *Lactobacillus* and several gram negative organisms relative to the Mock DNA control.

A native stool sample was used to determine the effect of DNA stabilization reagent on the overall microbial community composition. An additional element contributing to the differential community composition observed using the Mo Bio kit may be explained by analyses which showed that native stool samples were most dramatically affected by the presence of nucleotide stabilization reagent when extracted with Mo Bio. This indicates a potential incompatability of the Mo Bio kit with the DNA/RNA Shield stabilization reagent, which was also used to stabilize the commercially available Mock Microbial Community. This putative chemical incompatibility may have affected the microbial community composition observed in all DNA/RNA Shield-suspended samples extracted using the Mo Bio kit. This includes the whole cell mock community samples, which are available only suspended in DNA/RNA Shield. On the other hand, the increase in DNA yield per gram of stool in the presence of stabilization reagent used together with the Zymo Research kits is perhaps unsurprising because all components were manufactured by Zymo Research and were likely optimized to be used together. However, we have shown that the stabilization reagent can also be used successfully with the QIAamp kit. Although there is a decrease in yield, the reagent does not cause a decrease in the DNA quality. It should be noted, however, that our analyses show the use of DNA/RNA Shield, alters the observed abundance of numerous taxa compared to native stool and this should be taken into consideration when planning studies and comparing results from studies which differ in their use of stabilization reagent.

Development of best practices and standardized methods for microbiota analysis is critical for the advancement of research in many fields including personalized nutrition, ecology, and food science/safety. It will be necessary to perform similar experiments as new technologies are developed in order to make informed choices when determining which methods will provide the most accurate data.

## DECLARATIONS

### Ethics approval and consent to participate

The institutional review board of the University of California, Davis approved this study and all participants provided written informed consent (clinicaltrails.gov registration number NCT02298725).

### Consent for publication

Not applicapble

### Availability of data and material

All 16S rRNA sequences used in this analysis were deposited in the Qiita database (https://qiita.ucsd.edu) under study ID 11427 and in the European Nucleotide Archive (ENA) under accession number ERP104979.

### Competing interests

The authors declare that they have no competing interests.

### Funding

Research reported in this publication was supported in part by a grant from the National Dairy Council and the Campbell Soup Company and the USDA Agricultural Research Service project numbers 2032-51000-001-00D and and 2032-51530-022-00D.

### Authors' contributions

RH performed bioinformatics analyses and together with MEK wrote the main text of the manuscript. ZA performed DNA extraction, PCR, compilied data and contributed to writing the methods section. NK designed and managed the human study which provided stools for these analyses and provided editorial input for the manuscript.

## Acknowledgements

The authors would like to acknowledge Dr. Maria L. Marco for contributing support and ideas to this project. We would additionally like to thank Dr. Benjamin Callahan for his assistance in understanding DADA2 and diagnosing contaminant sequences in our dataset.

Mention of trade names or commercial products in this publication is solely for the purpose of providing specific information and does not imply recommendation or endorsement by the U.S. Department of Agriculture. USDA is an equal opportunity provider and employer.

## References

1. Manor, O. and E. Borenstein, Systematic Characterization and Analysis of the Taxonomic Drivers of Functional Shifts in the Human Microbiome. Cell Host Microbe, 2017. 21(2): p. 254–267.

2. Lloyd-Price, J., et al., Strains, functions and dynamics in the expanded Human Microbiome Project. Nature, 2017. 550(7674): p. 61–66.

3. Howard, M.M., T.H. Bell, and J. Kao-Kniffin, Soil microbiome transfer method affects microbiome composition, including dominant microorganisms, in a novel environment. FEMS Microbiol Lett, 2017. 364(11).

4. Oliverio, A.M., M.A. Bradford, and N. Fierer, Identifying the microbial taxa that consistently respond to soil warming across time and space. Glob Chang Biol, 2017. 23(5): p. 2117–2129.

5. Ward, C.S., et al., Annual community patterns are driven by seasonal switching between closely related marine bacteria. ISME J, 2017. 11(6): p. 1412–1422.

6. Neave, M.J., et al., Endozoicomonas genomes reveal functional adaptation and plasticity in bacterial strains symbiotically associated with diverse marine hosts. Sci Rep, 2017. 7: p. 40579.

7. Lemas, D.J., et al., Alterations in human milk leptin and insulin are associated with early changes in the infant intestinal microbiome. Am J Clin Nutr, 2016. 103(5): p. 1291–300.

8. Noble, E.E., et al., Early-Life Sugar Consumption Affects the Rat Microbiome Independently of Obesity. J Nutr, 2017. 147(1): p. 20–28.

9. Zieliska, S., et al., The choice of the DNA extraction method may influence the outcome ń of the soil microbial community structure analysis. MicrobiologyOpen, 2017.

10. Lozupone, C.A., et al., Meta-analyses of studies of the human microbiota. Genome Res, 2013. 23(10): p. 1704–14.

11. Aird, D., et al., Analyzing and minimizing PCR amplification bias in Illumina sequencing libraries. Genome biology, 2011. 12(2): p. R18.

12. Schirmer, M., et al., Insight into biases and sequencing errors for amplicon sequencing with the Illumina MiSeq platform. Nucleic acids research, 2015. 43(6): p. e37–e37.

13. Sinha, R., et al., Assessment of variation in microbial community amplicon sequencing by the Microbiome Quality Control (MBQC) project consortium. Nat Biotechnol, 2017.

14. Konstantinidis, K.T. and J.M. Tiedje, Genomic insights that advance the species definition for prokaryotes. Proceedings of the National Academy of Sciences of the United States of America, 2005. 102(7): p. 2567–2572.

15. Nguyen, N.-P., et al., A perspective on 16S rRNA operational taxonomic unit clustering using sequence similarity. npj Biofilms and Microbiomes, 2016. 2: p. 16004.

16. Schloss, P.D. and S.L. Westcott, Assessing and improving methods used in operational taxonomic unit-based approaches for 16S rRNA gene sequence analysis. Appl Environ Microbiol, 2011. 77(10): p. 3219–26.

17. Kopylova, E., et al., Open-Source Sequence Clustering Methods Improve the State Of the Art. mSystems, 2016. 1(1).

18. Tikhonov, M., R.W. Leach, and N.S. Wingreen, Interpreting 16S metagenomic data without clustering to achieve sub-OTU resolution. The ISME journal, 2015. 9(1): p. 68.

19. Edgar, R.C., UNOISE2: improved error-correction for Illumina 16S and ITS amplicon sequencing. bioRxiv, 2016: p. 081257.

20. Callahan, B.J., et al., DADA2: High-resolution sample inference from Illumina amplicon data. Nat Methods, 2016. 13(7): p. 581–3.

21. Amir, A., et al., Deblur Rapidly Resolves Single-Nucleotide Community Sequence Patterns. mSystems, 2017. 2(2).

22. Eren, A.M., et al., Minimum entropy decomposition: unsupervised oligotyping for sensitive partitioning of high-throughput marker gene sequences. ISME J, 2015. 9(4): p. 968–79.

23. Rosen, M.J., et al., Denoising PCR-amplified metagenome data. BMC bioinformatics, 2012. 13(1): p. 283.

24. Edgar, R.C., Accuracy of microbial community diversity estimated by closed- and open-reference OTUs. PeerJ, 2017. 5: p. e3889.

25. Callahan, B.J., P.J. McMurdie, and S.P. Holmes, Exact sequence variants should replace operational taxonomic units in marker gene data analysis. bioRxiv, 2017: p. 113597.

26. Vishnivetskaya, T.A., et al., Commercial DNA extraction kits impact observed microbial community composition in permafrost samples. FEMS microbiology ecology, 2014. 87(1): p. 217–230.

27. Bahl, M.I., A. Bergström, and T.R. Licht, Freezing fecal samples prior to DNA extraction affects the Firmicutes to Bacteroidetes ratio determined by downstream quantitative PCR analysis. FEMS microbiology letters, 2012. 329(2): p. 193–197.

28. Guo, F. and T. Zhang, Biases during DNA extraction of activated sludge samples revealed by high throughput sequencing. Applied microbiology and biotechnology, 2013. (10): p. 4607–4616.

29. Garcia-Mazcorro, J.F., et al., Influence of whole-wheat consumption on fecal microbial community structure of obese diabetic mice. PeerJ, 2016. 4: p. e1702.

30. Plante, C.J. and S. Stinson, Recolonization and cues for bacterial migration into mock¹ deposit-feeder fecal casts. Aquatic microbial ecology, 2003. 33(2): p. 107–115.

31. Kable, M.E., et al., The Core and Seasonal Microbiota of Raw Bovine Milk in Tanker Trucks and the Impact of Transfer to a Milk Processing Facility. MBio, 2016. 7(4).

32. Caporaso, J.G., et al., Global patterns of 16S rRNA diversity at a depth of millions of sequences per sample. Proceedings of the National Academy of Sciences, 2011. 108(Supplement 1): p. 4516–4522.

33. Hamady, M., et al., Error-correcting barcoded primers for pyrosequencing hundreds of samples in multiplex. Nature methods, 2008. 5(3): p. 235–237.

34. Bokulich, N.A., et al., Next-generation sequencing reveals significant bacterial diversity of botrytized wine. PloS one, 2012. 7(5): p. e36357.

35. Caporaso, J.G., et al., QIIME allows analysis of high-throughput community sequencing data. Nature methods, 2010. 7(5): p. 335–336.

36. Edgar, R.C., Search and clustering orders of magnitude faster than BLAST. Bioinformatics, 2010. 26(19): p. 2460–2461.

37. DeSantis, T.Z., et al., Greengenes, a chimera-checked 16S rRNA gene database and workbench compatible with ARB. Applied and environmental microbiology, 2006. 72(7): p. 5069–5072.

38. Segata, N., et al., Metagenomic biomarker discovery and explanation. Genome biology, 2011. 12(6): p. R60.

39. Li, K., et al., Analyses of the microbial diversity across the human microbiome. PloS one, 2012. 7(6): p. e32118.

40. Eren, A.M., et al., Oligotyping: differentiating between closely related microbial taxa using 16S rRNA gene data. Methods in Ecology and Evolution, 2013. 4(12): p. 1111–1119.

41. Leite, F.L., et al., Comparison of fecal DNA extraction kits for the detection of Mycobacterium avium subsp. paratuberculosis by polymerase chain reaction. Journal of veterinary diagnostic investigation, 2013. 25(1): p. 27–34.

42. Ruggieri, J., et al., Techniques for Nucleic Acid Purification from Plant, Animal, and Microbial Samples. Sample Preparation Techniques for Soil, Plant, and Animal Samples, 2016: p. 41–52.

43. Nechvatal, J.M., et al., Fecal collection, ambient preservation, and DNA extraction for PCR amplification of bacterial and human markers from human feces. Journal of microbiological methods, 2008. 72(2): p. 124–132.

